# Dynamic Changes in the Urinary Proteome of Normal Pregnant Mice Carrying Phenylketonuria-Affected Fetuses

**DOI:** 10.64898/2026.09.20.752939

**Authors:** Lili Guo, Youhe Gao

**Affiliations:** Gene Engineering Drug and Biotechnology Beijing Key Laboratory, College of Life Sciences, Beijing Normal University, Beijing 100871, China

**Keywords:** phenylketonuria, fetal origin, maternal urinary proteomics, biomarkers, early pregnancy screening, WGCNA, LASSO

## Abstract

Phenylketonuria (PKU) is an autosomal recessive disorder caused by *PAH* gene defects and can cause irreversible neurological damage as early as the fetal period; however, non-invasive prenatal monitoring methods are lacking. In this study, Pah^+/−^ female mice were mated with Pah^+/−^ male mice and wild-type male mice, respectively. Urine samples were collected longitudinally from pregnant mice at 10 time points during gestation from Day 1 to Day 19. Data-independent acquisition (DIA) quantitative proteomics combined with GO/KEGG analysis, WGCNA, and a dual-strategy LASSO-random forest approach was used to screen candidate biomarkers. The results showed that stable differentially expressed proteins between groups were detectable as early as Day 1, and the differences persisted throughout gestation and exhibited temporal dynamics. Permutation tests indicated that the probability of random generation of the differential protein combination was low (0.04–0.06), suggesting that it was not randomly generated. The intersection of the two strategies yielded five proteins—GINM1, PRG4, LY6A, NTM, and YIPF3—and after including GLO1, a total of six candidate biomarkers were obtained. The candidate proteins had interaction or co-expression associations with 14 proteins involved in PKU mechanisms, with GLO1 serving as a network hub. This study found that the maternal urine proteome can non-invasively reflect the systemic response induced by fetal-origin PKU, providing proof of concept and candidate biomarker resources for non-invasive early-pregnancy screening of PKU.

## 1. Introduction

Phenylketonuria (PKU) is an autosomal recessive inherited disorder of amino acid metabolism caused by PAH gene variants. It was first reported by the Norwegian physician Asbjørn Følling in 1934 ^[1]^. The core mechanism of the disease is the loss of phenylalanine hydroxylase activity, whereby phenylalanine cannot be normally converted to tyrosine, leading to substantial accumulation in the body and the generation of neurotoxic metabolites; if not promptly treated, it can cause irreversible neurological damage, with clinical symptoms mostly manifesting as intellectual disability and neurological deficits ^[2,3]^. PKU can occur in newborns, infants, children, and even adults, with the highest incidence among newborns. Early clinical manifestations are insidious and easily overlooked ^[3]^. The incidence of this disease varies across regions worldwide. The incidence among newborns in the United States is approximately 5.3–7.4 per 100,000 ^[4]^, and approximately 1 in 10,000 in Europe ^[5]^, imposing a heavy burden on affected families and public health.

In terms of pathogenesis, the core pathogenic basis of PKU is mutations in the *PAH* gene on chromosome 12, while a small number of cases are caused by mutations in genes involved in BH4 cofactor synthesis and metabolic pathways. These mutations lead to structural abnormalities and loss of activity of hepatic phenylalanine hydroxylase, which cannot catalyze the hydroxylation of phenylalanine (Phe) to tyrosine, resulting in persistent accumulation of blood Phe and its toxic metabolites phenylpyruvic acid, phenyllactic acid, and phenylacetic acid^[6,7]^. High intrauterine concentrations of Phe can cross the placenta and the fetal blood-brain barrier, competitively inhibiting the transport of essential amino acids such as tyrosine and tryptophan in the brain, directly causing deficient synthesis of key neurotransmitters such as dopamine and serotonin^[8,9]^. Meanwhile, a high-Phe environment can inhibit neuronal mitochondrial respiratory chain function, induce oxidative stress activation and massive ROS accumulation, and hinder fetal brain myelin development and neuronal proliferation and differentiation, ultimately leading to clinical phenotypes such as irreversible intellectual impairment, epilepsy, and motor developmental _delay_[9,10].

It is particularly important to distinguish maternal PKU, which refers to the mother herself being affected and intrauterine high Phe exposure causing fetal damage, from fetal-origin PKU, which refers to normal maternal phenylalanine metabolism, in which only the fetus develops metabolic abnormalities in utero due to PAH deficiency and triggers a maternal systemic response through the maternal-fetal interface. The latter does not involve maternal genetic variants and is an important source of current clinical missed diagnosis. At present, the domestic clinical PKU prevention and control system mainly relies on postnatal newborn screening and dietary intervention therapy, and still lacks mature prenatal functional diagnostic methods. The existing shortcomings are reflected in three aspects: (1) Newborn screening can only achieve postpartum diagnosis, but neurological damage has already occurred during the fetal stage, missing the intrauterine intervention window; (2) traditional genetic testing only detects mutation sites and cannot assess residual enzyme activity or the true metabolic phenotype; there is genotype-phenotype dissociation, making it difficult to distinguish mild and severe subtypes and to guide individualized early intervention; (3) it cannot identify secondary fetal-origin PKU injury caused by abnormal maternal-fetal interactions. Such cases do not involve maternal genetic variants and are an important source of clinical missed diagnosis. Therefore, developing non-invasive biomarkers for fetal PKU is a key breakthrough for achieving early screening, early diagnosis, and early intervention.

The Pah^−/−^knockout mouse model completely lacks PAH protein, can reproduce core pathological phenotypes of severe PKU such as hyperphenylalaninemia, neurotransmitter depletion, and myelin damage, and is widely used in preclinical PKU research^[10,14,15]^. In this study, pregnant mice with normal maternal phenotype (Pah^+/−^, normal phenotype) were used to carry Pah^−/−^homozygous affected offspring, specifically simulating the maternal-fetal interaction disturbance caused by fetal PAH deficiency. Two breeding schemes were established: the experimental group (DT) consisted of Pah^+/−^ females mated with Pah^+/−^ males, and the offspring could include PKU-affected fetal mice; the control group (CT) consisted of Pah^+/−^ females mated with wild-type males, and all offspring had a normal phenotype. Maternal urine from pregnant mice was used to capture the systemic molecular responses induced by pathological fetuses at the maternal interface and the dynamic changes in the urinary proteome during pregnancy.

## 2 Materials and Methods

### 2.1 Experimental Animals and Model Establishment

A total of 15 wild-type C57BL/6J male mice, 15 HE (*Pah*^+/−^) male mice, and 30 HE (*Pah*^+/−^) female mice, aged 10–12 weeks and of SPF grade, were selected. The mice were purchased from Shanghai Model Organisms Center, Inc. The animal license number was SCXK (Beijing) 2024-0003. All mice were housed in an SPF animal room at 22–25 °C and 50–60% humidity under a 12 h/12 h light/dark cycle, with free access to food and water. All experimental procedures were reviewed and approved by the Ethics Committee of the College of Life Sciences, Beijing Normal University, with the approval number CLS-AWEC-B-2022-003. After 1 week of acclimatization, the experiments were initiated.

Experimental Grouping: Normal pregnant mice carrying Pah-KO phenylketonuria (PKU) fetuses were assigned to the experimental group (Disease team, DT; normal maternal phenotype), whereas normal pregnant mice carrying normal fetuses were assigned to the control group (Control team, CT).

#### DT

*Pah*^+/−^ female mice × *Pah*^+/−^ male mice. The offspring may include *Pah*^+/+^ (WT), *Pah*^+/−^ (heterozygous, HE), and *Pah*^−/−^ (homozygous, HO) mice, with *Pah*^−/−^ mice being PKU fetuses.

#### CT

*Pah*^+/− female mice × *Pah*+/+ (WT) male mice. The offspring include only WT and HE mice, with no HO mice^_。_

Mating was performed daily at 18:00 by co-housing females and males at a female-to-male ratio of 2:1. Vaginal plugs were examined at 8:00 the following morning, and the day when a plug was observed was recorded as gestational day 0.5.

### 2.2 Urine Sample Collection

The sampling time points were Day 1, Day 3, Day 5, Day 7, Day 9, Day 11, Day 13, Day 15, Day 17, and Day 19. On each sampling day, from 20:00 to 8:00 the next morning, each pregnant mouse was placed individually in a separate metabolic cage overnight for urine collection; the urine volume collected from a single mouse per collection was ≥500 μL. After collection, urine samples were centrifuged at 4 °C and 3000 rpm for 10 min to remove cell debris and impurities. The supernatant was aliquoted into cryovials and stored at −80 °C, and repeated freeze–thaw cycles were strictly avoided.

### 2.3 Genotyping of Offspring Mice

Tail tissue was clipped from newborn mice and lysed overnight with proteinase K in a metal bath at 56 °C. Genomic DNA was obtained through DNA binding, washing, and elution. Three-primer PCR amplification was performed in a 20 μL total reaction system containing 0.5μL each of P1/P2/P3 primers, 10μL of 2× TransDirect Mouse Genotyping SuperMix, 2μL of template DNA, and 6.5μL of ddH2O. The PCR program was as follows: initial denaturation at 94 °C for 3 min; 34 cycles of 94 °C for 30s, 60 °C for 30s, and 72 °C for 30s; and final extension at 72 °C for 5min. PCR products were separated by 1% agarose gel electrophoresis at 120V for 30min and then imaged. Interpretation criteria were as follows: WT showed only a 380 bp band; HE showed two bands at 380 bp and 644 bp; HO showed only a single 644 bp band.

### 2.4 Sample Processing

#### Protein extraction

Urine samples were thawed at 4 °C and centrifuged at 12,000 g for 10 min at 4 °C. The supernatant was collected, and prechilled anhydrous ethanol was added at a volume ratio of 1:3, followed by precipitation at −20 °C for 12h. After centrifugation at 12,000g for 30 min at 4 °C, the supernatant was discarded. The pellet was dried and resuspended in lysis buffer, sonicated for 3 min, rotated at 4 °C for 2h, and the protein concentration was determined by the Bradford method.

#### Protein digestion

A total of 100μg protein was diluted to 200μL with NH_4_HCO_3_. DTT was added to a final concentration of 20 mM, and the mixture was reduced at 37 °C for 1h. After cooling, IAA was added to a final concentration of 50mM, and alkylation was performed at room temperature in the dark for 30 min. Buffer exchange was performed using a 10 kDa ultrafiltration tube, followed by two washes with UA buffer and three washes with NH_4_HCO_3_. Trypsin was added at an enzyme-to-protein mass ratio of 1:50, and digestion was performed at 37 °C overnight. The peptide filtrate was collected by centrifugation at 14,000g. Peptides were desalted using an HLB solid-phase extraction cartridge and stored at−20°C after lyophilization.

### 2.5 LC-MS/MS Mass Spectrometry Analysis

The lyophilized peptides were dissolved in 0.1% formic acid, and peptide concentration was determined using a BCA kit. All samples were diluted to 0.5 μg/μL. A 10 μL aliquot of the diluted sample was centrifuged at 12,000 g and 4 °C for 30 min, and the supernatant was collected and spiked with 1μL IRT. Proteomic analysis was performed in 96-well plates. Three intra-plate QCs were set up per plate, prepared by mixing equal amounts of 10 randomly selected digested samples from that plate; meanwhile, a pooled quality control sample, QCmix, was prepared as an equal mixture of all digested samples, and these were used for intra-plate and overall experimental quality control, respectively.

The entire detection platform consisted of an UltiMate 3000 nano-liquid chromatography system coupled with an Orbitrap Exploris 480 high-resolution mass spectrometer. An integrated C18 capillary column (75μm inner diameter, 50cm length) was used, with the column temperature maintained at 60 °C. The elution flow rate was 1.5 μL/min, and the linear gradient elution program with acetonitrile (containing 0.1% formic acid) was as follows: the organic phase proportion increased from 5% to 20% within 15.5 min, from 20% to 30% within 5 min, to 50% within 1 min, rapidly to 90% within 0.1 min and maintained for 1.3 min, and finally decreased to 5% within 0.1 min, followed by equilibration for 2 min. Mass spectrometry data were acquired in positive ion mode using data-independent acquisition (DIA): the full MS scan resolution was 120,000, the AGC target was 300%, the maximum ion injection time was 50 ms, and the scan mass-to-charge ratio range was 350–1200 m/z; the DIA MS2 scan resolution was 30,000, the AGC target was 200%, the maximum ion injection time was 50 ms, and the higher-energy collisional dissociation (HCD) collision energy was 30%.

### 2.6 Data Searching

The raw DIA data were imported into Spectronaut v19 software and searched against the Swiss-Prot mouse database. Trypsin/P was used for digestion, with up to 2 missed cleavages allowed. The peptide length was set to 7–52 amino acids. Carbamidomethylation of cysteine was set as a fixed modification. Variable modifications included protein N-terminal acetylation and methionine oxidation. The false discovery rate (FDR) was ≤1% at both the peptide and protein levels. Protein quantification abundance was calculated as the sum of the fragment ion peak areas of all matched peptides.

### 2.7 Data Analysis

Missing value filtering was performed on the protein quantification matrix of all urine samples covering the ten gestational time points from Day 1 to Day 19: proteins with a missing rate >50% in both groups were removed; among the missing values of the retained proteins, those with a missing rate >50% in a single group were filled with half of the minimum value detected for that protein within that group, and the remaining missing values were imputed using the KNN algorithm. The criteria for screening differentially expressed proteins were as follows: fold change (FC) ≥1.5 or ≤0.67 and two-tailed unpaired t-test P value <0.05. In addition, group-specific protein screening rules were established: proteins with a missing rate >50% in both groups were excluded;□a detection rate ≥50% in at least one group;□≤2 samples detected in one group and a detection rate >50% in the other group. Such specific proteins are difficult to identify by conventional differential analysis and were separately included in the subsequent analysis workflow.

### 2.8 Bioinformatics and Statistical Analysis

#### Functional enrichment analysis

GO/KEGG functional enrichment of differentially expressed proteins was performed using the DAVID database and visualized with SRplot, and the top 15 enriched terms were extracted for each time point.

#### WGCNA analysis

The log2-transformed protein quantification matrix of the full dataset was used to construct a scale-free co-expression network with scale-free R² > 0.8. Hierarchical clustering combined with dynamic tree cutting was used to divide modules. Module eigengenes (MEs) were calculated and subjected to Pearson correlation analysis with gestational time and fetal genotype.

#### Biomarker screening

Two complementary screening strategies were used: early-stage-specific screening (using 64 samples from Day1/Day3/Day5) and cross-time-point universal screening (using all 202 samples). Both employed an intersection strategy of LASSO-penalized logistic regression (5-fold stratified cross-validation, binomial deviance scoring, lambda.1se criterion) and random forest (n_estimators = 500) to reduce false positives associated with a single method. LASSO regression shrinks the coefficients of irrelevant features to zero via an L1 penalty and has been widely used in biomarker screening of high-dimensional omics data, including the construction of urine metabolite- and gene-based diagnostic models for pregnancy diseases such as preeclampsia ^[16,17]^.

#### Statistical software and visualization

Statistical analyses were performed using R software and Python 3. Principal component analysis (PCA) and partial least squares discriminant analysis (PLS-DA) were used for sample pattern recognition and quality control. Protein abundance trend clustering analysis, sample correlation heatmaps, protein abundance trend plots, biomarker–mechanism correlation heatmaps, and STRING interaction network plots were generated using tools such as ggplot2, ComplexHeatmap, and matplotlib. P < 0.05 was considered statistically significant.

## 3. Results

### 3.1 Genotyping Results of Offspring

Genotypes of the offspring mice were identified by 1% agarose gel electrophoresis (Figure 1). The amplified fragment for WT was 380 bp; HE showed both 380 bp and 644 bp bands; and HO showed only a 644 bp band. The results showed that WT, HE, and HO genotypes were all detectable among the offspring of the experimental group, whereas only WT and HE were detected among the offspring of the control group, with no HO individuals observed. The electrophoretic band sizes completely matched the genotyping criteria, indicating that the breeding system in this study can stably produce PKU offspring and corresponding controls, thereby providing a reliable animal model foundation for urinary proteomics research on fetal-origin phenylketonuria.

**Figure 1.**
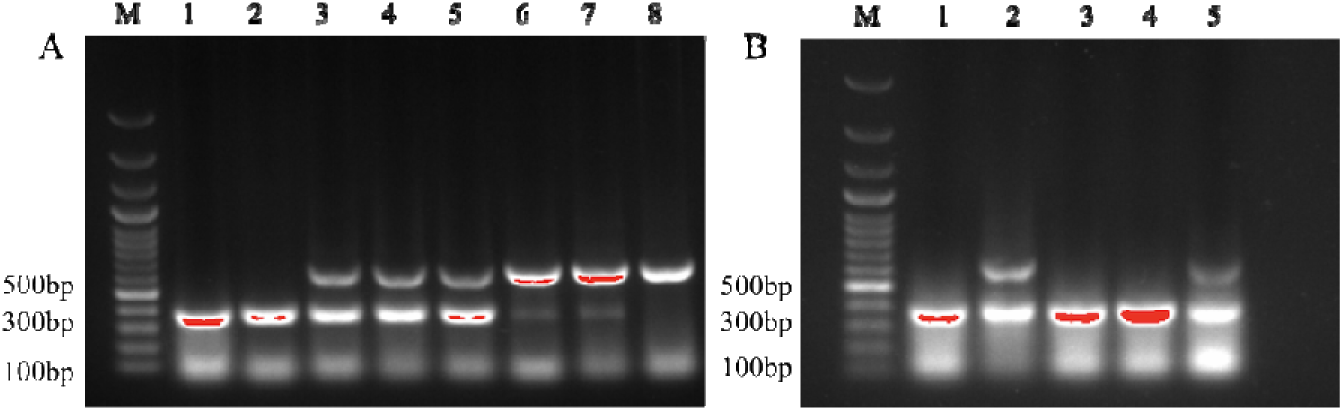
PCR genotyping of the *Pah* gene in offspring mice detected by 1% agarose gel electrophoresis. M: DNA Marker; (A) Offspring from the experimental group: bands 1/2 are WT, bands 3–5 are HE, and bands 6–8 are HO; (B) Offspring from the control group: bands 1/3/4 are WT, and bands 2/5 are HE.

### 3.2 Phenotype Identification Results of Offspring

The gross phenotype and body weight of offspring mice of different genotypes were recorded (Figure 2). In terms of appearance, WT and HE mice had black coat color, whereas HO mice exhibited markedly lighter gray coat color, which is associated with impaired melanin synthesis. Regarding body weight, male WT and HE mice showed steady growth from 6 weeks of age, with similar body weights between the two groups, whereas HO homozygotes exhibited growth retardation, and their body weight was significantly lower than that of the WT and HE groups. The above phenotypes of coat color lightening and growth retardation are consistent with the characteristic manifestations of PKU model mice, further supporting the successful establishment of the experimental model.

**Figure 2.**
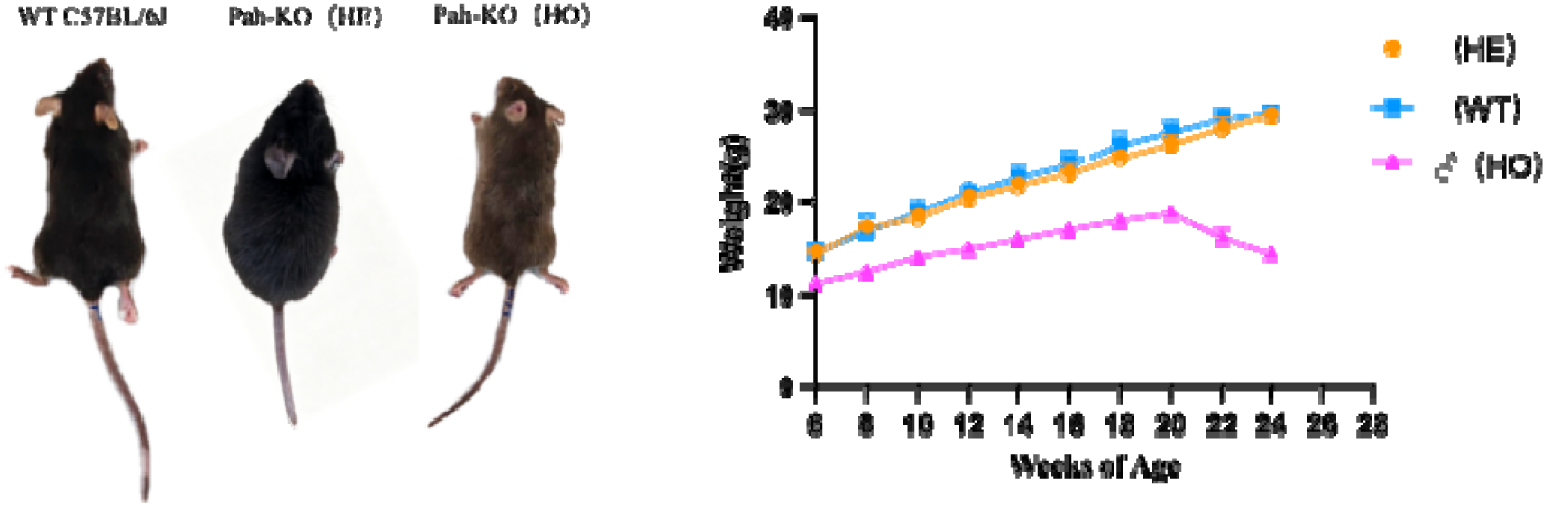
Gross phenotype and body weight growth curve of Pah-KO mice. Left panel: WT and HE mice had similar appearance with black coat color, whereas HO mice exhibited coat color lightening. Right panel: Body weight change curves of male mice from 6 to 24 weeks of age; WT and HE mice showed normal weight gain, whereas HO homozygous knockout mice exhibited growth retardation, with body weight significantly lower than that of the WT and HE groups.

### 3.3 Proteomic Results

#### 3.3.1 Model Construction and Sample Collection

The pregnancy rates and total gestation duration of the two groups of pregnant mice showed no significant differences (P > 0.05). In the control group, 12 mating pairs were successfully mated, and in the experimental group, 10 mating pairs were successfully mated. No miscarriage or death of dams occurred during the experimental period.

#### 3.3.2 Overall Changes in the Urinary Proteome During Pregnancy

In this study, a total of 207 urine samples were collected from pregnant mice in the experimental group (n = 10) and the control group (n = 12) during gestation (Day 1–19). Across all samples (Table 1 and Table S1, Figure 1A), a total of 2,591 proteins were identified (FDR < 1.0%), with an average of 1,128 proteins per sample. After quality control screening, 202 samples were used for subsequent analysis (16 samples were excluded due to insufficient protein concentration).

**Table 1.** Urine protein identification result.

| Group | Number of Samples | Time Point | Number of Identified Proteins |
| --- | --- | --- | --- |
| DT | 104 | Day1~Day19 | 1174 ± 295 |
| CT | 114 | Day1~Day19 | 1090 ± 286 |
| All | 218 | Day1~Day19 | 1128 ± 292 |

#### 3.3.3 Quality Control of the Mass Spectrometry Platform

To evaluate the technical reproducibility of the mass spectrometry-based proteomics experiments, the coefficient of variation (CV) of protein signal intensities in intra-plate quality control (QC) samples was calculated. As shown in Figure 3, the median protein-level CVs of the intra-plate QC samples QC1–QC3 were 31.2%, 48.5%, and 15.7%, respectively, and the CVs of the vast majority of proteins were below 50%. The Pearson correlation coefficients among QCmix replicate samples ranged from 0.89 to 0.97 (Table S2). These results demonstrate that the technical reproducibility of the entire DIA proteomic detection system is reliable.

**Figure 3.**
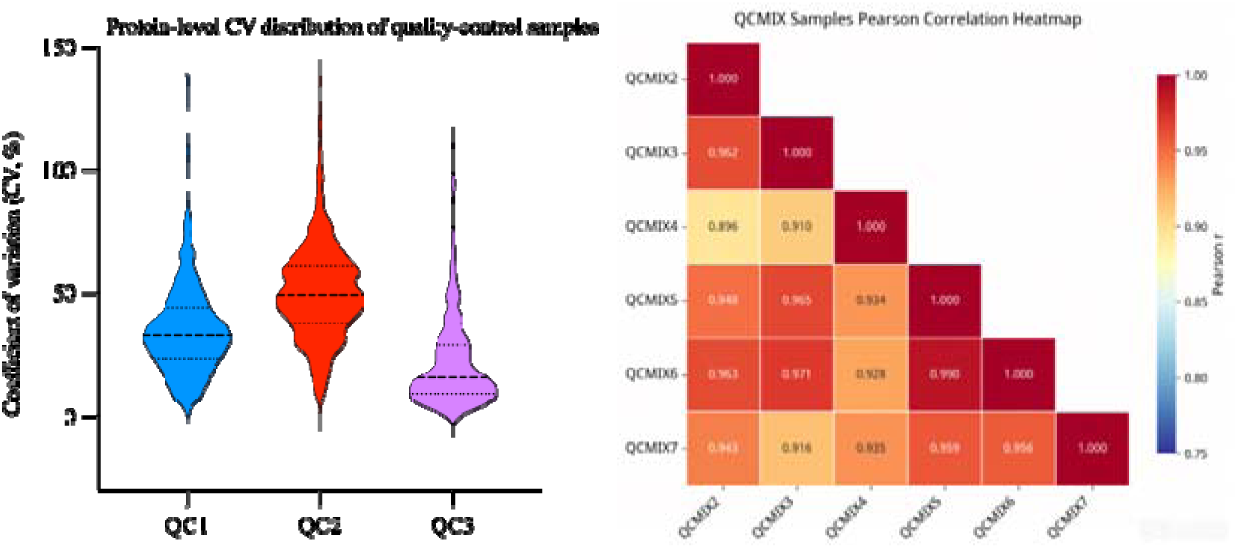
Distribution of protein-level coefficient of variation (CV) in QC samples. Violin plots show the distribution of coefficient of variation (CV, %) of protein signal intensities for the three intra-plate QC samples (QC1–QC3). The solid black line inside each violin indicates the median CV, and the dashed lines denote the 25th and 75th quartiles. Lower CV values indicate better technical reproducibility of proteomic measurement.

#### 3.3.4 Permutation Test to Verify the Reliability of Differential Proteins

To rule out random artifacts, we performed random combination simulations at each time point (Day 1–19). For each time point, we randomly selected the same number (n) of proteins from the m differential proteins to form random combinations, and calculated the random generation probability—the probability that a random combination performed as well as or better than the observed combination. A lower probability indicates a lower likelihood of random generation and stronger biological relevance. As shown in Table 2, 9 of the 10 time points (Day 1, 3, 5, 7, 11, 13, 15, 17, and 19) had very low random generation probabilities (0.04–0.06), suggesting that the differential protein combinations at most time points were not randomly generated and had statistical significance and potential biological relevance.

**Table 2.**
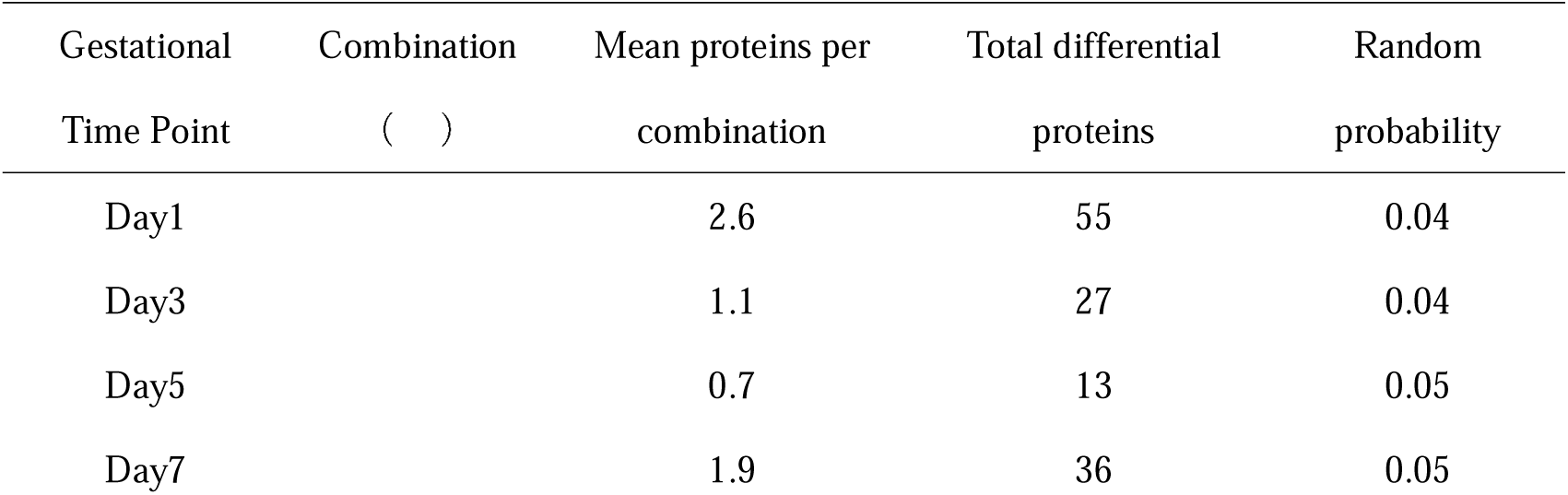
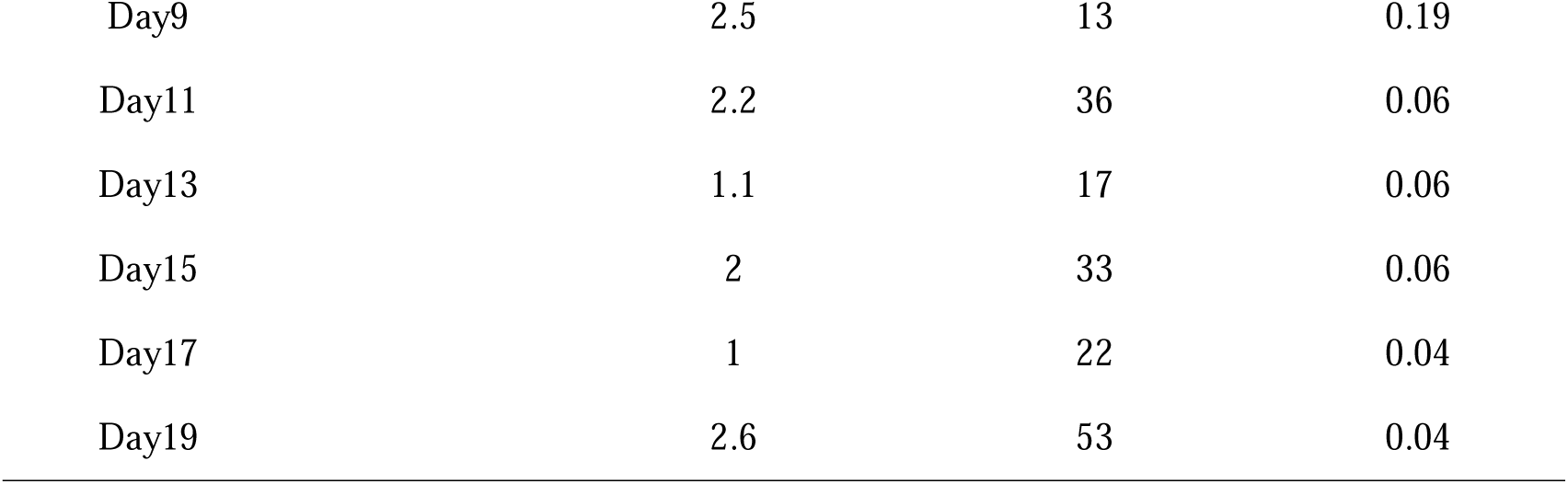
Validation of the non-randomness of DEPS combinations across ten gestational time points.

#### 3.3.5 Distribution of Differential Proteins at Different Gestational Time Points

The dynamic differential distribution of the urinary proteome during pregnancy between mice carrying PKU offspring and those carrying normal offspring is shown in Figure 4. At each time point, the numbers of up- and down-regulated differential proteins exhibited strong temporal fluctuations: early pregnancy was dominated by down-regulated proteins; as pregnancy progressed, the number of down-regulated proteins gradually decreased; after Day 9, up-regulated proteins predominated, and the number of up-regulated proteins peaked at Day 19. UpSet intersection analysis suggested that most differential proteins were single-time-point-specific changes, with only a few proteins commonly differentially expressed across multiple gestational stages. This indicates that the maternal perturbation caused by fetal-origin PKU is highly temporally dynamic, and maternal response patterns differ markedly across different gestational weeks.

**Figure 4.**
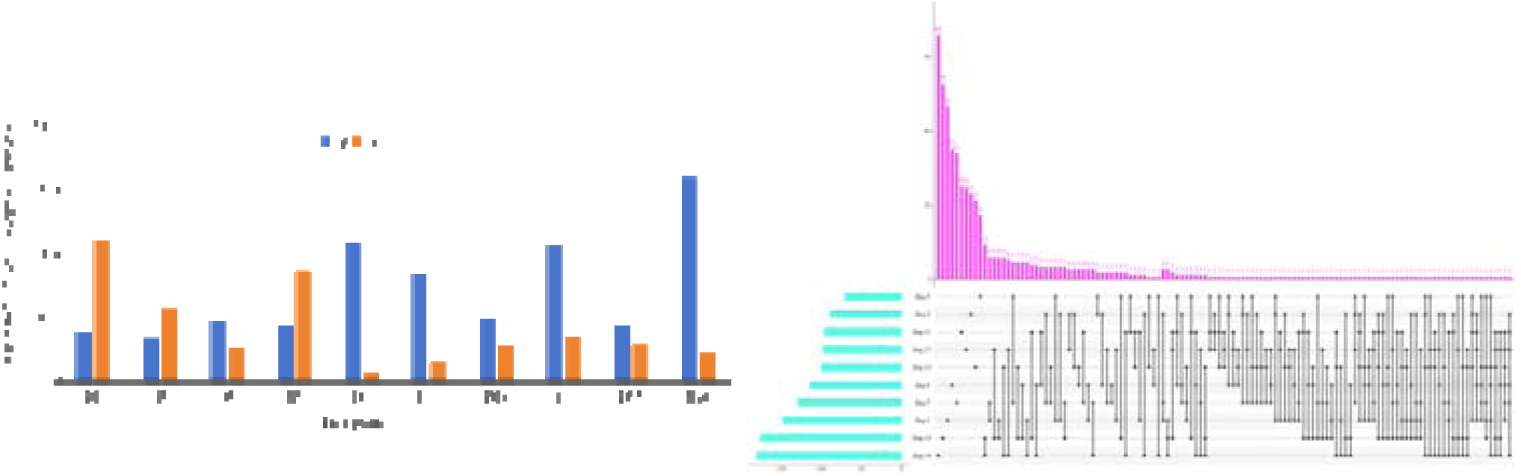
Distribution characteristics of differential proteins in maternal urine from gestational Day 1 to Day 19: (A) Bar chart of the number of differential proteins at each gestational week: blue represents significantly up-regulated proteins, and orange represents significantly down-regulated proteins; (B) UpSet plot of differential protein intersections: the horizontal bars on the left indicate the total number of differential proteins for a single gestational week, the vertical bars at the top indicate the total number of proteins in the corresponding intersection set, and the dot-line connections represent the differential proteins shared by the corresponding combinations of gestational weeks.

#### 3.3.6 Temporal Detection Characteristics of PKU Differential Proteins

Differential proteins at a total of 10 gestational time points in pregnant mice carrying PKU offspring relative to those carrying normal offspring (Figure 5, Table S3). Figure 5A shows 45 key PKU proteins (9 directly related + 36 strongly related). Early pregnancy was dominated by pterin metabolism and amino acid transport: on Day 1, PCBD1 (pterin-4α-carbinolamine dehydratase), a key enzyme in the BH4 regeneration cycle, was detected, along with significantly up-regulated TTR^[18,19]^; on Day 3, the amino acid transporter SLC6A14, which uses L-phenylalanine as a substrate, was detected^[20]^; on Day 5, metabolic enzymes such as the GST family, GGCT, and SHMT1 were predominantly detected[21]; from Day 7 onward, the glyoxalase/glutathione detoxification system (GLO1, HAGH/GLO2) and mitochondrial respiratory chain proteins (UQCRC1, COX5B) appeared as a group, suggesting activation of the methylglyoxal–AGE–neurotoxicity axis and mitochondrial oxidative stress^[22]^; on Day 9, the directly causative PKU protein phenylalanine hydroxylase (PAH) was detected for the first time, accompanied by concentrated activation of ubiquitin–proteasome/ERAD components^[6,23]^; Day 11 was the time point with the densest antioxidant network; aromatic L-amino acid decarboxylase (AADC/DDC), GCLM, SOD2, and NQO1 were detected simultaneously, reflecting coupling between neurotransmitter synthesis and oxidative stress defense^[21,24]^; Day 13 was characterized by a mitochondrial energy metabolism cluster (ACADVL, UQCRC1, COX5B) and FSTL1; on Day 15, the myelin lysosomal lipid metabolism enzymes GALC and ARSA and the folate–methionine cycle enzymes MTR and MTHFD1 appeared in a concentrated manner, consistent with myelin formation defects and one-carbon metabolism disorders in PKU^[25–27]^; on Day 17, the endoplasmic reticulum glycoprotein quality control enzyme UGGT1 was detected^[6]^; on Day 19, PAH was detected again, and the pterin family bifunctional protein Gephyrin and the PLP synthase PNPO appeared; the product of PNPO, PLP, is precisely an essential cofactor of AADC, forming a cross-time-point evidence chain of “vitamin B6–neurotransmitter synthesis”^[28–30]^. Cross-stage stable proteins such as KIAA0319L, ECM1, and PRG4 can serve as priority candidates for temporal biomarkers. Figure 5B,C show 23 proteins stably and specifically detected between groups at ≥3 time points (e.g., NUA, C9, ECM1, PRG4), which are difficult to capture by conventional differential analysis; their functions involve oxidative stress, immune inflammation, myelin damage, neurodevelopment, and protein homeostasis, highly consistent with the core pathology _of PKU_[9,31].

**Figure 5.**
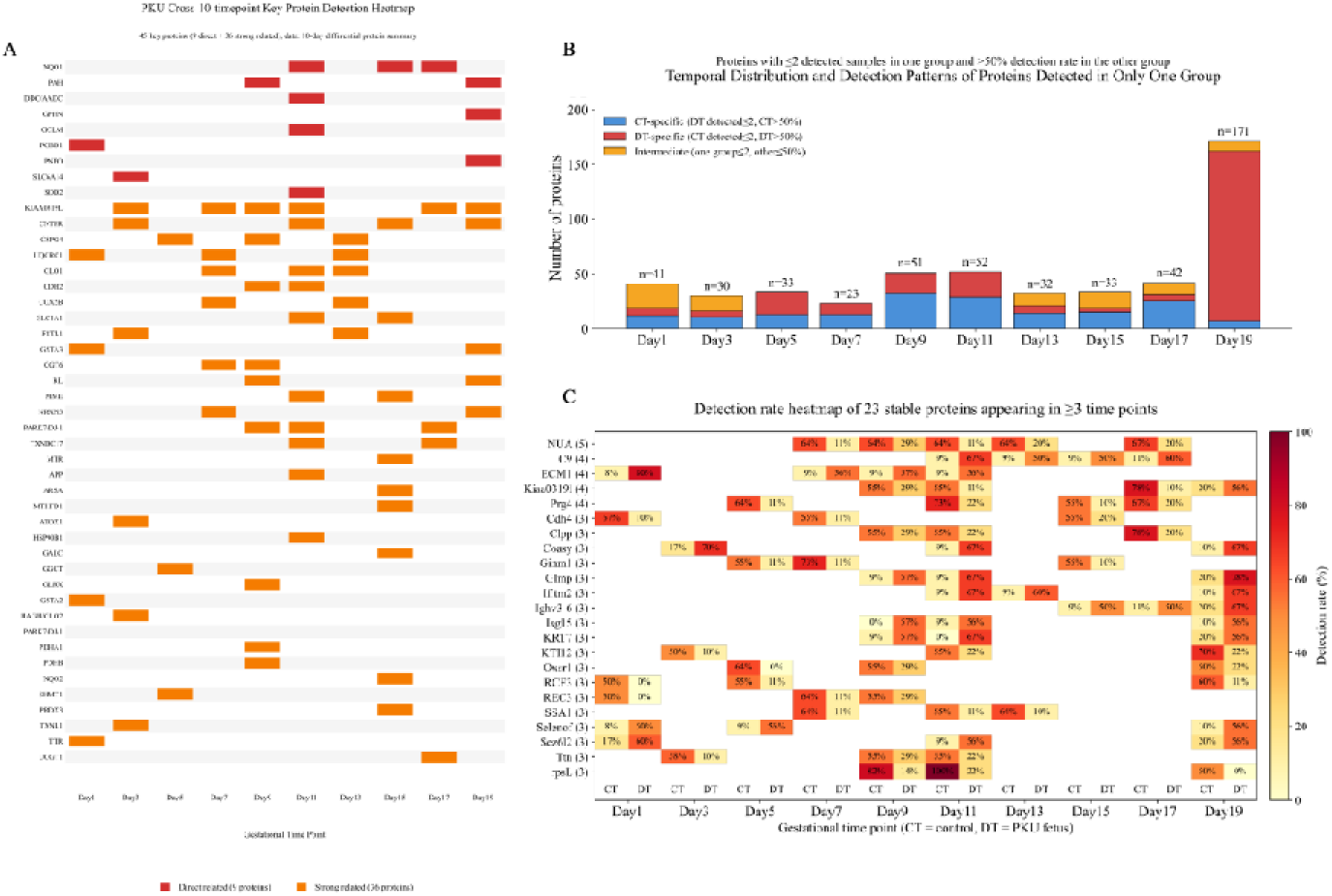
Heatmap of key PKU-associated proteins detected across ten time points and temporal distribution plus detection-rate heatmap of proteins detected exclusively in one group. (A) Detection profiles of 45 key proteins (9 directly related proteins in red, 36 strongly related proteins in orange) from Day 1 to Day 19, ranked by relevance. (B) Stacked bar charts across the ten time points (blue = CT-specific, red = DT-specific, orange = intermediate type). (C) Detection-rate heatmap of the 23 stable proteins identified in ≥3 time points for CT and DT groups. CT: dams carrying normal fetuses; DT: dams carrying PKU-affected fetuses.

#### 3.3.7 WGCNA Co-expression Network Analysis Reveals the Maternal Systemic Metabolic Response to Fetal-Origin PKU

To systematically dissect the maternal systemic response induced by fetuses with phenylketonuria (PKU) in normal dams, weighted gene co-expression network analysis (WGCNA) was performed on 1,247 log2-transformed proteins based on urinary proteomic data at 10 time points during gestation^[32,33]^ (table S4). A total of five modules were identified (Figure 6A): brown (149), turquoise (678), yellow (148), blue (229), and green (24); 19 proteins were not included in subsequent analysis. Module–phenotype association analysis (Figure 6B) showed that brown was significantly positively correlated with DT (r = 0.708, p = 7.7 × 10□³²), and turquoise was elevated in DT (r = 0.278, p = 6.4×10□□); yellow (r = −0.483, p = 3.6×10□¹³) and blue (r = −0.267, p = 1.3 × 10□□) were correlated with CT; green showed no significant association (r = −0.068, p = 0.34). No module was significantly associated with gestational days (p =0.294–0.781, |r| ≤ 0.074), suggesting that the co-expression changes were mainly driven by PKU fetal mice rather than by the progression of pregnancy itself. Module eigengenes (Figure 6C) further confirmed that the brown module was enriched in oxidative stress/metabolic proteins such as SOD2, GLO1, NQO1, Klotho, and SHMT1, which were continuously highly expressed throughout gestation in DT; the turquoise module was enriched in DDC, GSTM2, GCLM, ARSA, PCBD1, etc., and maintained at relatively high levels in DT; the yellow (PRG4, FSTL1, SLPI, etc.) and blue (TTR) modules were highly abundant in CT. GO-BP temporal enrichment analysis (Figure 6D) showed that oxidative stress–glutathione, synaptic signaling, immune response, and protein degradation pathways were gradually enriched in mid-to-late pregnancy, fully recapitulating the pathological processes of maternal oxidative stress, mitochondrial disorder, and protein homeostasis imbalance induced by hyperphenylalaninemic fetuses^[34–36]^. Module members encompassed functional clusters including oxidative stress defense, amino acid metabolism, and membrane transport, indicating that the maternal response triggered by fetal-origin PKU is a systematic network remodeling coordinated by multiple proteins^[33,37]^. These results indicate that the urinary proteome can non-invasively reflect the metabolic status of the maternal–fetal interface during pregnancy.

**Figure 6.**
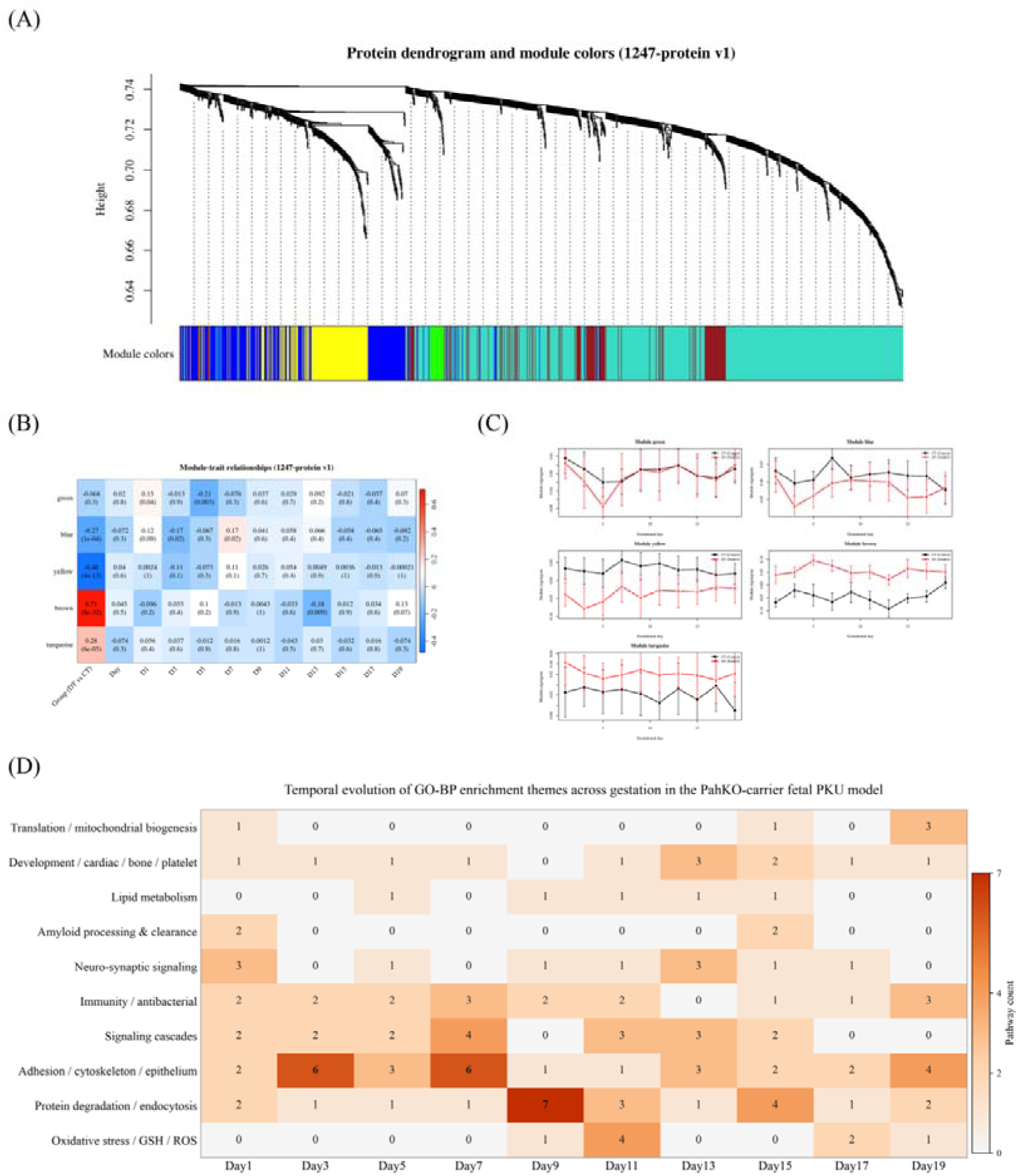
Weighted gene co-expression network analysis (WGCNA) and GO enrichment analysis of the urinary proteome. (A) Protein dendrogram and module color annotation: a scale-free co-expression network was constructed using all proteins in the full matrix. (B) Module–phenotype relationship heatmap: each cell shows the Pearson correlation coefficient (r) and significance p value (in parentheses) between the module eigengene (ME, the first principal component of proteins within the module) and phenotypes . Red indicates positive correlation, and blue indicates negative correlation. (C) Temporal changes in module eigengenes across gestational days: data points are group means, and error bars indicate 95% confidence intervals. (D) Temporal evolution heatmap of GO-BP enrichment themes of differential proteins in the PKU model.

#### 3.3.8. KEGG Pathway and Protein–Protein Interaction Network Reveal a Coordinated Phenylalanine–One-Carbon Metabolism–Oxidative Stress Response

To comprehensively evaluate the differential characteristics of the gestational urinary proteome between the DT and CT groups, multidimensional integrative analysis was performed based on differential proteins from the 10 time points. PCA showed that the two groups could be roughly distinguished, and supervised PLS-DA achieved complete separation of the two groups (LOO accuracy 99.0%, Figure7A). The sample correlation heatmap showed high intra-group correlation and marked inter-group separation, with clear hierarchical clustering according to gestational day (Figure 7B).

**Figure 7.**
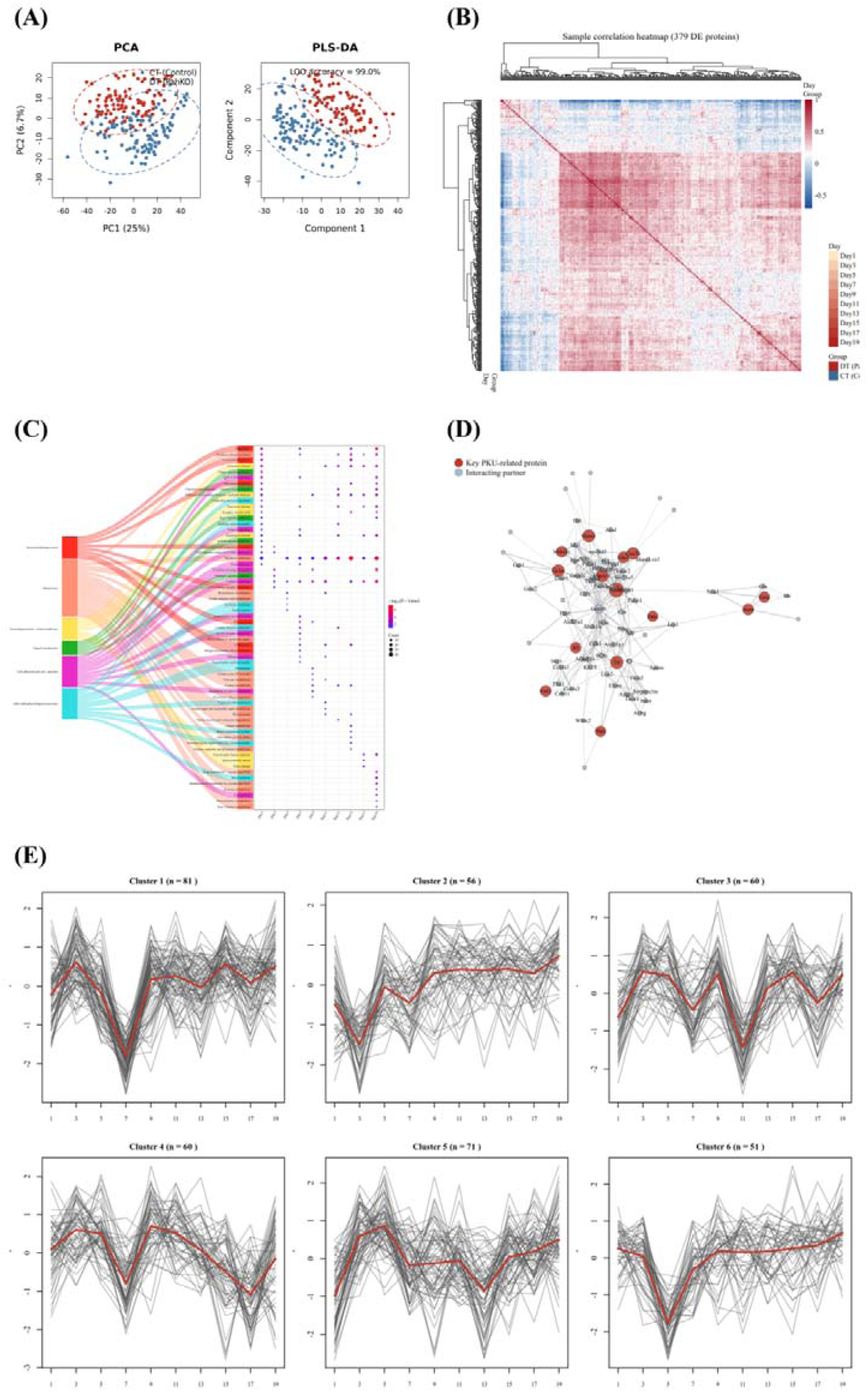
Overall compositional analysis of DEPs in the gestational urinary proteome of DT vs. CT. A: PCA and PLS-DA (CT vs. DT; PLS-DA LOO accuracy 99.0%). B: Sample correlation heatmap based on differential proteins from ten time points (annotated by Day and Group). C: KEGG pathway enrichment Sankey diagram–time scatter plot. D: STRING PPI core network (red: PKU-related key proteins; gray: interaction partners; edge thickness confidence). E: Mfuzz trend clustering (6 clusters; red thick line indicates the cluster mean).

KEGG enrichment showed that differential proteins were mainly enriched in metabolism, immune phagocytosis, neurodegenerative pathways, signal transduction, and cell adhesion. Phenylalanine metabolism and tyrosine metabolism pathways were significantly enriched in mid-to-late pregnancy, corroborating each other with the repeated detection of PAH on Day 9 and Day 19 and the temporal detection of the amino acid transporter SLC6A14 in early pregnancy, suggesting that the maternal metabolic response triggered by fetal PKU metabolic abnormalities gradually amplifies with advancing gestation^[38,39]^. ROS and oxidative phosphorylation pathways were significantly enriched, consistent with the brown and turquoise modules in WGCNA; these two modules contain core proteins for ROS scavenging and glyoxalase detoxification, such as SOD2, GLO1, and GCLM^[40–42]^. ARSA is related to myelin lipid homeostasis, and its enrichment peak on Day 11 coincided with the GO window of “response to oxidative stress, positive regulation of ROS metabolism,” suggesting coordinated initiation of antioxidant defense and myelin lipid metabolism in mid-pregnancy^[43]^. Synaptic vesicle cycle and neurodegenerative pathways such as Alzheimer’s, Parkinson’s, and Huntington’s diseases were significantly enriched, linking with the detection of AADC/DDC on Day 11 and the “vitamin B6–neurotransmitter synthesis” evidence chain, suggesting that maternal urine can capture fetal neurometabolic perturbations. In the STRING protein–protein interaction network (confidence score ≥ 0.4, Figure 7D), 14 key PKU differential proteins (SOD2, TTR, SHMT1, UQCRC1, GCLM, COX5B, MTHFD1, GLO1, ARSA, FSTL1, KL, GALC, SLPI, DDC, etc.) served as the core and formed interaction modules with proteins related to phenylalanine hydroxylation, one-carbon/folate metabolism, pterin metabolism, oxidative stress, neurotransmitter synthesis, and myelin lipid metabolism, suggesting that fetal PKU drives a coordinated maternal metabolic–oxidative stress–protein homeostasis network response^[44,45]^. Mfuzz (Figure 7E) classified the differential proteins into 6 trend clusters. For example, Cluster 4 (peak at D9, containing TTR) and Cluster 6 (valley at D5, containing PCBD1) corresponded to thyroxine transport and pterin regeneration, respectively, reflecting pathway activation windows at different gestational weeks and fully recapitulating the stage-specific maternal response to the metabolic burden of PKU fetuses^[46]^. In summary, PKU fetuses can leave multidimensional, dynamic, and discriminable proteomic imprints in maternal urine, and proteins related to metabolic remodeling, oxidative stress, and neurotransmitter synthesis constitute the core response module^[8,36,47,48]^.

### 3.9 Dual-Strategy Screening of Candidate Biomarkers Using LASSO and Random Forest

To systematically screen early diagnostic biomarkers in the maternal urine of pregnant mice with fetal-origin PKU, we adopted two strategies—early-stage (Day1–Day5, n=64) and cross-time-point (Day1–Day19)—and performed combined screening using LASSO and random forest. Early-stage-specific screening: LASSO (lambda.1se =3.393) selected 25 candidate proteins, and their intersection with the random forest Top50 yielded 12 early core proteins, as shown in Figure 8, among which Ginm1 (Q91WR6) ranked first in both algorithms. This model achieved an AUC of 1.000 in early pregnancy (Day1–Day5) and 0.844–0.960 in mid-pregnancy, but decreased in late pregnancy, suggesting clear early-window specificity.

**Figure 8.**
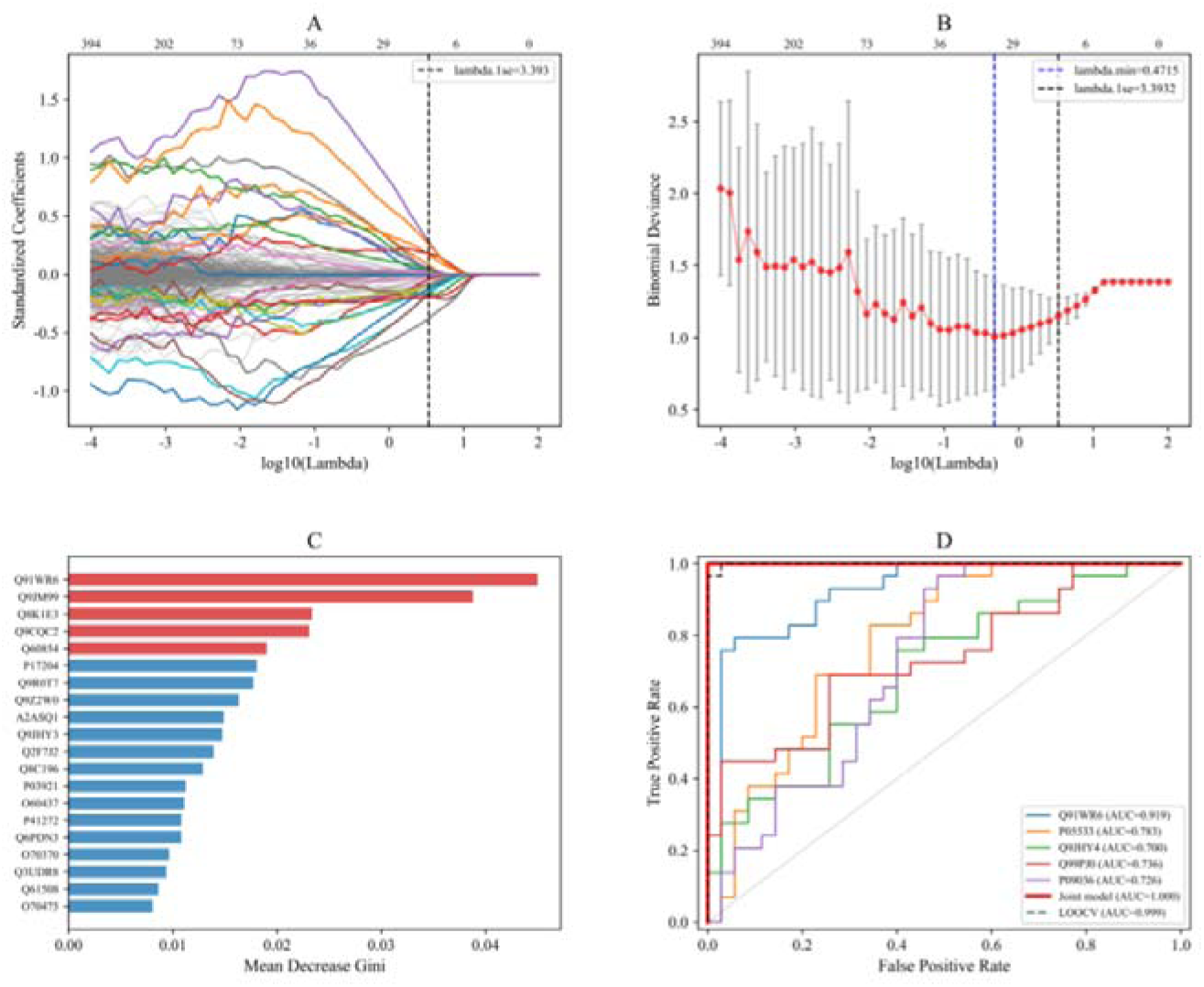
(A–D) Early-stage-specific screening: (A) LASSO coefficient trajectory plot; the black dashed line indicates lambda.1se (3.393, 25 non-zero coefficients); (B) LASSO 5-fold cross-validation plot, with red points representing the mean binomial deviance, error bars representing standard error, the blue dashed line indicating lambda.min, and the black dashed line indicating lambda.1se; (C) random forest variable importance bar chart (Top 20), with the top 5 highlighted in red and the x-axis showing Mean Decrease Gini; (D) ROC curves, with colored thin lines representing univariate models of the top 5 LASSO coefficients, the thick red line representing the combined model of 25 candidate proteins (AUC = 1.000), and the black dashed line representing LOOCV (AUC = 0.999).

The cross-time-point screening results are shown in Figure 9. LASSO selected 38 candidate proteins at lambda.1se=0.0373, and their intersection with the random forest Top50 yielded 15 cross-time-point core proteins. The two strategies produced 5 overlapping proteins: Ginm1, Prg4, Ly6a, Ntm, and Yipf3, indicating that these proteins are robust biomarkers independent of the screening strategy. Among them, Ginm1 is the candidate biomarker with the greatest translational potential. Notably, Ntm (neurotrimin), whose knockout mice exhibit emotional learning deficits and anxiety-like behaviors, suggesting an important role in neurodevelopment and cognitive function^[49,50]^, was down-regulated in the urine of pregnant mice carrying PKU fetuses, possibly indirectly reflecting the impact of abnormal fetal neurodevelopment on the maternal microenvironment. Glo1 (glyoxalase 1) has the most direct association with the pathological mechanism of PKU. It is not only a core protein in the full-matrix screening but also a differential protein. It is a key rate-limiting enzyme of the glyoxalase system, catalyzing the conversion of the hemithioacetal formed by methylglyoxal (MG) and glutathione to S-D-lactoylglutathione, and is a core enzyme in cellular defense against MG toxicity^[51]^. Elevated MG levels are closely associated with neurodegenerative diseases ^[52]^. The up-regulation of Glo1 reflects a compensatory maternal response to fetal-derived methylglyoxal neurotoxicity, consistent with the grouped activation of the glyoxalase/glutathione system starting from Day 7. Ultimately, Glo1 and the 5 overlapping proteins were considered candidate biomarkers.

**Figure 9.**
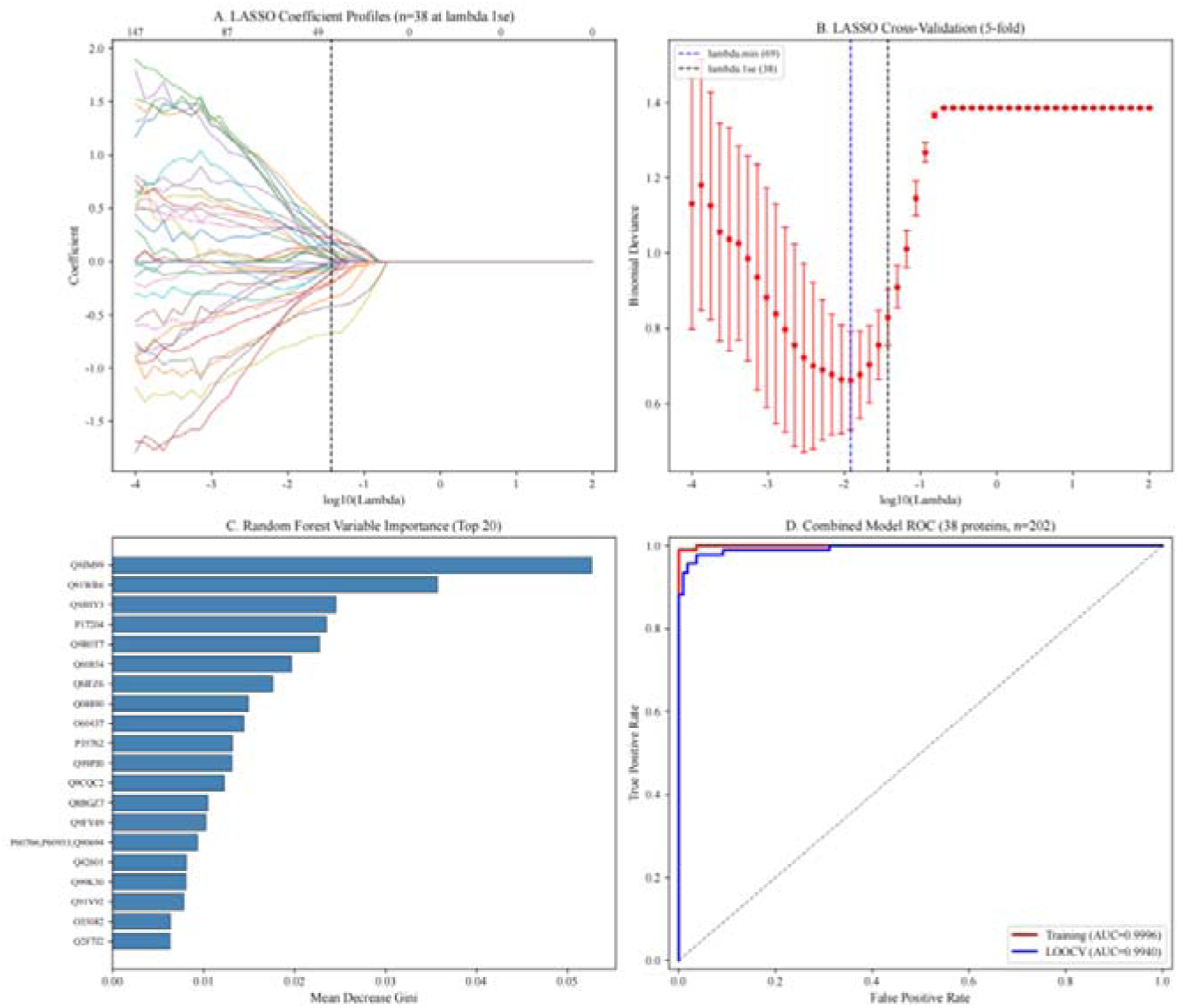
LASSO–random forest diagnostic biomarker screening based on full-matrix proteins, cross-time-point universal screening (Day 1–Day 19, n = 202): (A) LASSO coefficient trajectory plot, lambda.1se = 0.0373 (38 non-zero coefficients); (B) LASSO 5-fold cross-validation plot; (C) random forest variable importance Top 20; (D) ROC curves, combined model AUC = 0.9996, LOOCV AUC = 0.9940.

**Figure 10.**
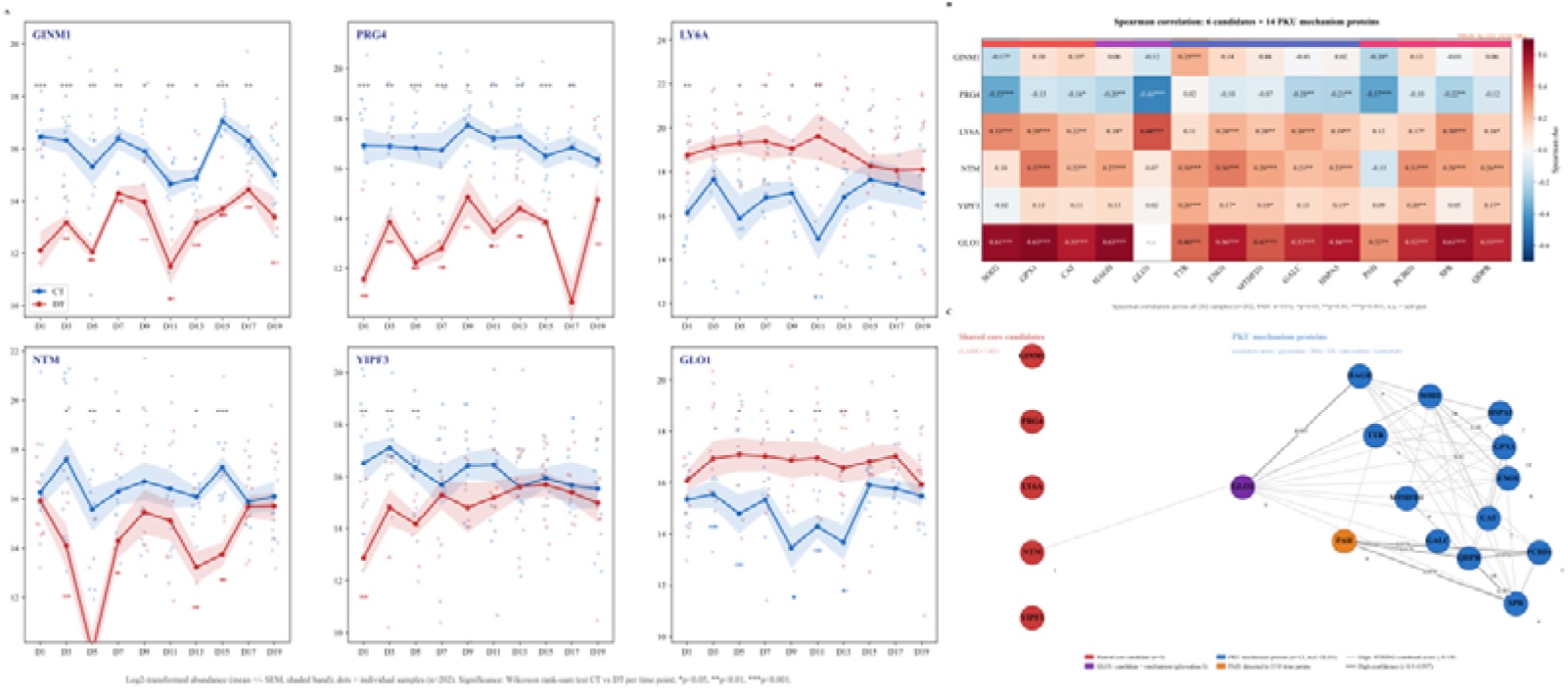
Abundance trends, expression correlations, and protein–protein interaction network validation of candidate biomarkers with PKU mechanism proteins. (A) Urinary abundance trends (log2 intensity) of the six candidate biomarkers across ten gestational time points from Day 1 to Day 19; the shaded area represents the mean ± confidence interval. Between-group comparisons were performed using the Wilcoxon rank-sum test with FDR correction (P < 0.05, P < 0.01, *P < 0.001). (B) Spearman expression correlation heatmap of the six candidate proteins and 14 PKU mechanism proteins. Color represents the Spearman correlation coefficient (red, positive correlation; blue, negative correlation); P < 0.05. (C) STRING protein–protein interaction network of the six candidate proteins (red nodes) and 14 PKU mechanism proteins (blue nodes) (Mus musculus).

#### 3.3.10. Validation of the Association Between Candidate Biomarkers and PKU Mechanism Proteins

To elucidate the association between the candidate biomarkers and the pathogenesis of PKU, the six candidate biomarkers and 14 PKU mechanism proteins obtained from full-matrix screening (oxidative stress: SOD2, GPX1, CAT; glyoxalase/carbonyl detoxification: HAGH, GLO1; BH4–phenylalanine metabolism: PAH, PCBD1, SPR, QDPR; neurometabolism/endoplasmic reticulum stress: TTR, ENO1, MTHFD1, GALC, HSPA5) were subjected to STRING interaction and Spearman correlation analyses. The STRING network showed that GLO1 was a hub node (degree 9), simultaneously connecting the glyoxalase detoxification, oxidative stress, and BH4 metabolic pathways; GLO1 was significantly positively correlated with the vast majority of mechanism proteins (r =0.32–0.63). The remaining five candidate proteins lacked direct physical interactions with the mechanism proteins and were mainly characterized by co-expression changes and indirect regulatory relationships. These results suggest that the urinary proteome of pregnant mice carrying PKU fetuses captures the oxidative stress, carbonyl stress, and BH4 metabolic responses transmitted from fetal metabolic disturbances through the maternal–placental interface, supporting the establishment of “candidate biomarker–PKU mechanism” evidence chain.

## 4. Discussion

This study is the first to use urinary proteomics to find that pregnant mice carrying PKU fetal mice showed disease-related differential signals, such as PCBD1 and TTR, as early as Day 1, and these signals persisted throughout the entire gestational period. Permutation tests showed that the random generation probability of these differential protein combinations was low (0.04–0.06), suggesting that they were not random noise but rather an early maternal response with potential biological significance. At Day 1, mouse embryos are still at the zygote-to-cleavage stage; why could detectable changes already appear in the maternal urinary proteome? This may be related to the following mechanisms. First, the embryonic genotype is already determined at fertilization. The DT group can produce *Pah*^−/−^ embryos, whereas the CT group does not produce *Pah*^−/−^ embryos; therefore, the two groups differ in embryonic genotype from the earliest stage. Second, preimplantation embryos can affect the maternal local microenvironment through the secretion of metabolites, proteins, extracellular vesicles, or small RNAs via oviductal and uterine fluids, and can even enter the maternal circulation to trigger a systemic response^[53–55]^. *Pah*^−/−^ embryos may exhibit abnormal phenylalanine metabolism due to PAH deficiency, leading to altered metabolite profiles, which are recognized by the mother and trigger adjustments in pterin metabolism, oxidative stress, or transport-related pathways. In this study, PCBD1 and TTR detected on Day 1 are involved in the BH4 regeneration cycle and thyroxine/retinol transport, respectively ^[56,57]^, suggesting that systemic changes at the metabolic and transport levels occur in the mother as early as the very early stage of pregnancy.

Functional enrichment showed that the maternal response gradually shifted from pterin–amino acid transport in early pregnancy to oxidative stress and myelin metabolic remodeling in mid-to-late pregnancy. WGCNA revealed that the maternal response was organized into modules involving oxidative stress defense, amino acid metabolism, and membrane transport, indicating a systemic network remodeling coordinated by multiple proteins^[58,59]^. KEGG and STRING analyses further revealed coordination among BH4 metabolism, one-carbon metabolism, and the antioxidant/glyoxalase system, forming a pathological cascade of “phenylalanine accumulation → one-carbon metabolism/BH4 imbalance → enhanced oxidative stress → impaired neurotransmitter synthesis.” Dual machine learning screening identified six candidate biomarkers: GINM1, PRG4, LY6A, NTM, YIPF3, and GLO1. GLO1 occupies a hub position in the interaction network and directly participates in the methylglyoxal–AGE neurotoxicity detoxification pathway; the remaining five proteins are mainly characterized by expression co-association, and their mechanisms in maternal–fetal interaction still require functional experimental validation. The six proteins showed significant between-group differences as early as the first trimester and weakened as pregnancy progressed, consistent with the goal of early screening. GLO1 has dual roles as both a candidate biomarker and a mechanism protein, supporting the central role of the methylglyoxal–AGE–neurotoxicity axis ^[60,61]^.

Clinical PKU mainly relies on newborn screening, and non-invasive intrauterine detection methods are lacking. This study, at the mouse model level, found that fetal inherited metabolic diseases can leave detectable temporal proteomic imprints in maternal urine, providing proof of concept and candidate biomarkers for non-invasive early-pregnancy screening of PKU. Limitations include: limited sample size; low urinary protein abundance and large individual variability, with missing value imputation potentially introducing bias; no simultaneous detection of maternal blood, placenta, or fetal tissues; candidate biomarkers have not yet been validated in pregnant women; and the early-stage model AUC approached 1.000, requiring caution against overfitting and external validation in independent cohorts.

## 5. Conclusions

Differential proteomic analysis of maternal urine throughout gestation revealed regular, reproducible, and dynamic proteomic changes in pregnant mice carrying PKU fetuses. Permutation tests showed low random generation probabilities (0.04–0.06) at 9 of 10 time points (0.19 on Day9), indicating that the differential protein combinations were non-random and statistically significant. Functional enrichment revealed a temporal response shifting from pterin metabolism/amino acid transport in early pregnancy to oxidative stress and myelin remodeling in mid-to-late pregnancy. WGCNA showed modular organization of the maternal response, and KEGG and STRING analyses revealed coordinated phenylalanine–one-carbon metabolism–oxidative stress response. Dual-strategy screening identified five core candidates—GINM1, PRG4, LY6A, NTM, and YIPF3—with both early specificity and cross-time-point stability; including GLO1 yielded six candidate biomarkers, and candidate–mechanism associations were established. These findings provide proof of concept and a candidate biomarker basis for non-invasive early-pregnancy screening of PKU based on maternal urinary proteomics.

## Acknowledgements

This work was supported by the National Key Research and Development Program of China (Grant No. 2023YFA1801900) and the Beijing Natural Science Foundation (Grant Nos. L2604022, L246002). We thank all lab members for the assistance of animal feeding, urine sample collection and mass spectrometry data analysis.

## Author Contributions

Conceptualization: Y.G.; writing—review and editing: Y.G.; supervision: Y.G.; methodology: L.G.; formal analysis: L.G.; data curation: L.G.; writing—original draft preparation: L.G. All authors have read and agreed to the published version of the manuscript.

## Institutional Review Board Statement

The animal study protocol was approved by the Ethics Committee of the College of Life Sciences, Beijing Normal University, with the approval number CLS-AWEC-B-2022-003.

## Informed Consent Statement

Not applicable.

